# Molecular genetics of GLUT1DS Italian pediatric cohort: 10 novel related-disease variants and structural analysis

**DOI:** 10.1101/2022.09.26.509448

**Authors:** Alessia Mauri, Alessandra Duse, Giacomo Palm, Roberto Previtali, Stefania Maria Bova, Sara Olivotto, Sara Benedetti, Francesca Coscia, Pierangelo Veggiotti, Cristina Cereda

**Affiliations:** Department of Biomedical and Clinical Sciences, University of Milan, Milan, Italy; Newborn Screening and Genetic Metabolic Diseases Unit, V. Buzzi Children’s Hospital, Milan, Italy; Pediatric Neurology Unit, V. Buzzi Children’s Hospital, Milan, Italy; Structural biology center, Human Technopole, Milan, Italy

## Abstract

GLUT1 deficiency syndrome (GLUT1DS1; OMIM #606777) is a rare genetic metabolic disease, characterized by infantile-onset epileptic encephalopathy, global developmental delay, progressive microcephaly and movement disorders (e.g. spasticity and dystonia). It is caused by heterozygous mutations in the *SLC2A1* gene, which encodes the GLUT1 protein, a glucose transporter across the blood-brain barrier (BBB). Most commonly these variants arise *de novo* resulting in sporadic cases, although several familial cases with AD inheritance pattern have been described.

Twenty-seven Italian pediatric patients clinically suspect of GLUT1DS from both sporadic and familial cases have been enrolled.

We detected by trios sequencing analysis 25 different variants causing GLUT1DS. Of these, 40% of the identified variants (10 out of 25) had never been reported before, including missense, frameshift and splice site variants. Their X-ray structure analyses strongly suggested the potential pathogenic effects of these novel disease-related mutations, broadening the genotypic spectrum heterogeneity found in the *SLC2A1* gene. Moreover, 24% is located in a vulnerable region of the GLUT1 protein that involves transmembrane 4 and 5 helices encoded by exon 4, confirming a mutational hotspot in the *SLC2A1* gene. Lastly, we investigated possible correlations between mutation type and clinical and biochemical data observed in our GLUT1DS cohort, revealing that splice site and frameshift variants are related to a more severe phenotype and low CSF parameters.

**Author summary:** We investigated the molecular data of 27 pediatric patients clinically suspect of GLUT1 deficiency syndrome. By performing trios sequencing analysis, we highlighted ten novel disease-related variants, and their X-ray structure analyses, suggesting the pathogenic effects of these identified mutations. Moreover, the wide clinical and genetic heterogeneity observed in our cohort allowed possible correlations between mutation type and clinical and biochemical data. This analysis enabled to delineate that splice site and frameshift variants are related to a more severe phenotype and low CSF/glucose values. Further clinical and genetic/epigenetics studies could clearly the high phenotypic variability observed in these patients.

## Introduction

Glucose transporter-1 (GLUT1) deficiency syndrome (GLUT1DS), also known as De Vivo disease, is a rare genetic metabolic condition characterized by infantile-onset epileptic encephalopathy, global developmental delay, drug-resistant seizures, progressive microcephaly, and movement disorders (e.g. spasticity, and dystonia) (GLUT1DS1; OMIM #606777) [1–3]. Although the infantile-onset epileptic encephalopathy phenotype is the most reported presentation (~90% of affected individuals), some individuals develop atypical or mild phenotype (~10%), characterized by infantile-onset paroxysmal exercise-induced dyskinesia (PED) with or without seizures (GLUT1DS2; OMIM #612126) [4–6]. Other paroxysmal events, which may be triggered by prolonged exercise, fasting, or emotional stress, include sleep disturbance (e.g. sleep apnea), abnormal eye-head movements, and weakness/paralysis [7–10]. Low cerebrospinal fluid (CSF) glucose values (termed hypoglycorrhachia) in the absence of hypoglycemia, in combination with low-normal or abnormally low CSF lactate values represent the metabolic hallmark of GLUT1DS [11,12]. These patients are treated with ketogenic diet therapies (KDT) that provide an alternative fuel source for brain energy metabolism^12^. Effectiveness of the KDT has been observed by Leen *et al*., not only in patients with drug-resistant epilepsy, but also in patients with movement disorders without epilepsy [5,13–15]

The epidemiological data of GLUT1DS are limited and only about 600 cases have been reported in medical literature since 1991 [1]. An incidence of 1.65-2.22 per 100.000 births was recently estimated [16].

GLUT1DS is caused by heterozygous mutations (or rarely, homozygous mutations) in the solute carrier family 2 member 1 gene (*SLC2A1*, 1p34.2) [17–19]. The *SLC2A1* gene encodes the glucose transporter-1 (GLUT1). It belongs to the GLUT protein superfamily, composed of 14 members (GLUT1-14) facilitative membrane transporters, involved in the regulation of glucose delivery and metabolism [20]. All GLUT transporters share a common protein structure characterized by 12 α-helices transmembrane (TMH) domains, separated by a large intracellular (IC) loop between helices 6 and 7. The GLUT1 protein is primarily expressed in erythrocytes, in endothelial cells comprising the blood-brain barrier (BBB) and in astrocytes. GLUT1 facilitates glucose diffusion from the bloodstream across the BBB and astrocyte plasma membrane to the central nervous system, representing the fundamental vehicle by which glucose enters the brain [12]. Consequently, a genetic defect in this transporter would impair glucose supply to the brain, resulting in a cerebral energy deficiency that is likely accounting for the clinical manifestations of the GLUT1DS [17,21]. A 3D model for GLUT1 protein has been proposed, predicting a central aqueous channel communicating the intra- and extracellular environment with many amino acid residues crucial for glucose transport located around this central channel [22,23].

So far, a total of 240 pathogenic *SLC2A1* variants have been reported in GLUT1DS patients including deletions, missense, nonsense, frameshift, and splice site mutations (ClinicalStatistics.Varsome.com) [5,24,25]. Most commonly these mutations arise *de novo* resulting in sporadic cases, although several familial cases with autosomal dominant inheritance pattern have been reported [25,26].

All these observed mutations are considered to lead to GLUT1DS by a loss of GLUT1 function. Nonsense, frameshift and splice-sites *SLC2A1* variations resulting in 50% loss of GLUT1 protein are often associated with the early-onset classical phenotype, whereas mild or moderate forms of the disease are most likely associated with missense *SLC2A1* mutations resulting in 50-70% residual function of GLUT1 [5,15,24,27]. However, genotype-phenotype correlation remained elusive with high interindividual phenotypic variability, even between mutated members of the same family [15,24].

The purpose of the study was to investigate the molecular data of a large Italian cohort of 27 pediatric patients clinically suspect of GLUT1DS from both sporadic and familial cases; specifically, we performed trios sequencing analysis, highlighted ten novel disease-related mutations and their pathogenic effects by X-ray structure analyses, and then investigated possible correlations between mutation type and clinical and biochemical data.

## Results

### Clinical and biochemical features

The main clinical and biochemical characteristics of 27 patients are shown in Table 1. The most frequent phenotype (n = 23; 85,2%) represented the epilepsy. Among these 23 patients, 17 had an onset of seizures before the age of two years (early-onset classical phenotype; 73,9%), while in 6 patients the onset of seizures occurred later in life (late-onset classical phenotype; 22,7%). Seizure semiology was diverse (tonic-clonic, dyscognitive, epileptic spasms), and also the frequency varied from daily to sporadic. Intellectual disability (ID) ranged from mild (n = 4; 23,5%) to severe (n = 7; 41,1%) in patients with the early-onset classical phenotype (2 patients had moderate ID, one borderline ID and 2 normal ID, while data of one patient were unavailable); in patients with the late-onset classical phenotype, severity of ID ranged from mild (n = 2; 33,3%) to severe (n = 3; 50%) (one patient had normal ID). Movement disorders (ataxia, dystonia, choreoathetosis) are shown in 73,9% (n = 17) out of 23 of the patients with seizures (13 patients with the early-onset and 4 in late-onset classical phenotype). Microcephaly (head circumference less than 3rd percentile for age and sex) occurred in 1 of 17 patients with the early-onset classical phenotype (patient 3 and patient 14). Most of these patients did not respond to antiepileptic drugs, such as phenobarbital, and therefore they were treated with KDT (n = 19; 70,3%).

**Table 1:**
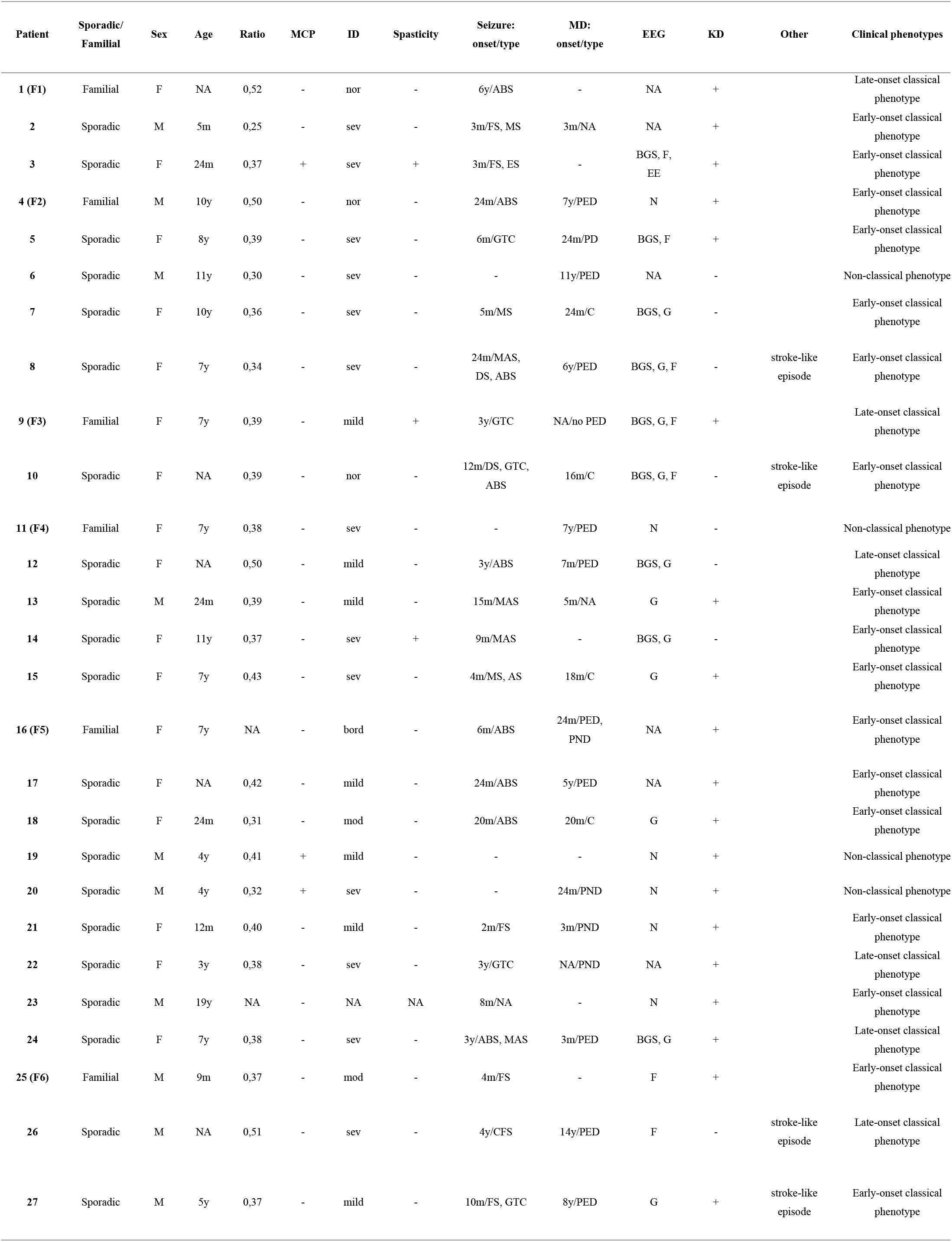
Clinical and biochemical data observed in the GLUT1DS Italian patients. Abbreviations: F, female; M, male; Age, age of diagnosis; Ratio, CSF/blood glucose ratio; NA, not available; +, yes; -, no; MCP, microcephaly; m, months; y, years; sev, severe; mod, moderate; ABS, absence seizure; CFS, complex febrile seizures; DS, dyscognitive seizures; EE, epileptic encephalopathy; ES, epileptic spasms; FS, focal seizures; GTC, generalized tonic-clonic seizures; MAS, myoclonic absence seizures; MS, myoclonic seizures; MD, movement disorder; C, chronic; PED, paroxysmal exertion-induced dyskinesia; PND, paroxysmal non exertion-induced dyskinesia; F, focal discharges; G, generalized discharges; N, normal; BGS, slow background slowing; KD, ketogenic diet.

Furthermore, the non-classical phenotype characterized by paroxysmal exercise-induced dyskinesia without seizures occurred in 3 out of 27 patients (11,1%). One of these 3 patients presented microcephaly (patient 20).

Finally, patient 19 presented only minimal symptoms: mild intellectual disability and microcephaly.

CSF/blood glucose ratios (reference range 0.6 mg/dL) in all patients ranged from 0.25 to 0.52 (mean 0.39; SD 0.06) (data of patient 16 and patient 23 were unavailable).

### Molecular data

Sequencing analysis of the 27 patients detected 25 different variants in the *SLC2A1* gene (Fig. 1a), including 15 missense (60%), 6 frameshift (24%), one nonsense (4%), one splice site (4%), one in-frame (4%) and one noncoding (4%) (Table 2). Of these, 10 mutations (40%) have never been described before. Moreover, 24% (6 out of 25 identified variants) is located on transmembrane helix 4 (TMH4) and transmembrane helix 5 (TMH5) encoded by exon 4, confirming a mutational hotspot in the *SLC2A1* gene observed by Pascual *et al*. [28].

**Table 2:**
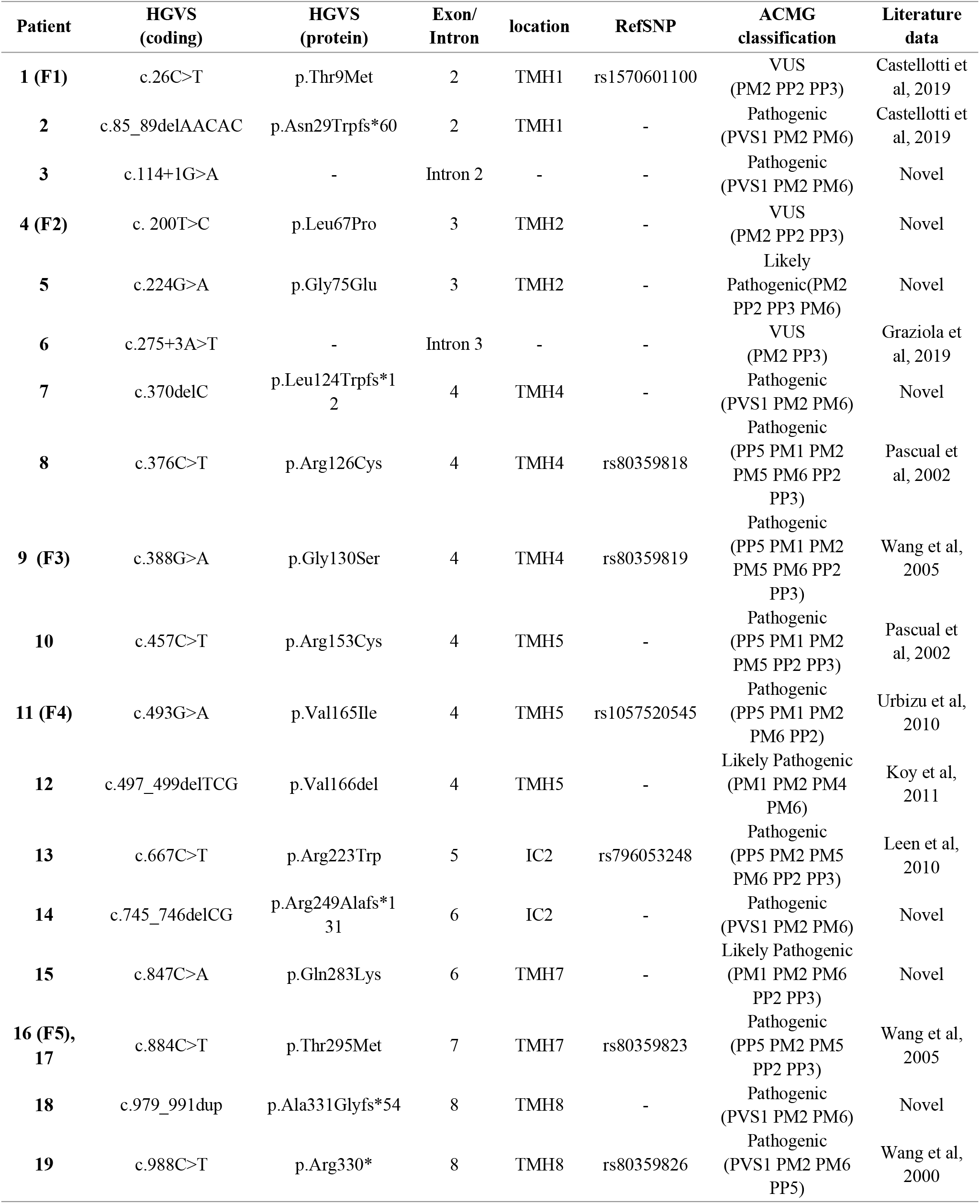
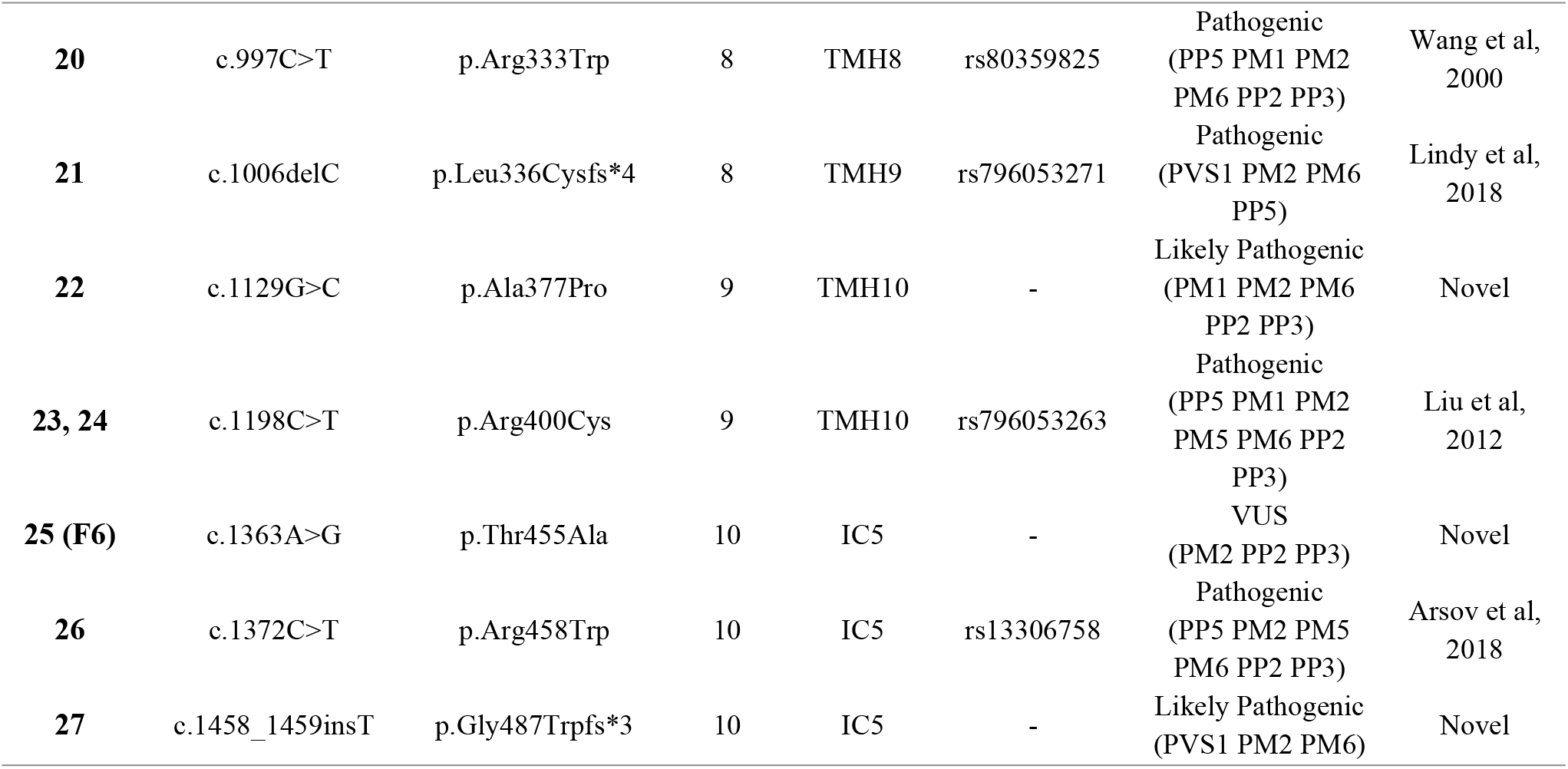
*SLC2A1* variants identified in 27 GLUT1DS patients with details on HGVS mutation, exon, location, refSNP, ACMG classification and literature reference for variants already reported. Abbreviations: refSNP, reference SNP; TMH, α-helices transmembrane; IC intracellular domain; VUS, variant of uncertain significance.

**Figure 1:**
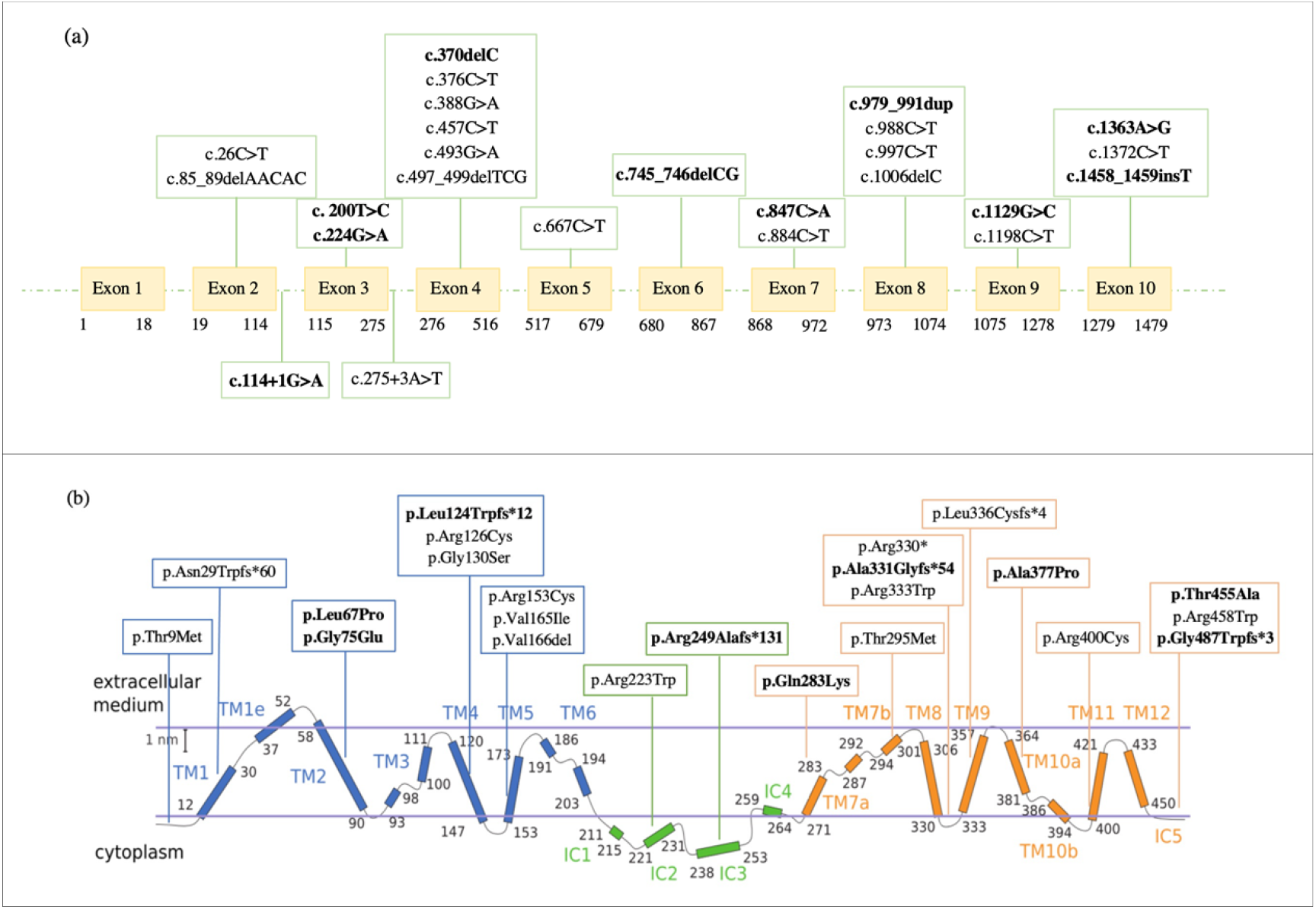
**a**, Distribution of 25 different variants in the *SLC2A1* gene identified in 27 patients with GLUT1DS. **b**, Representation of the GLUT1 protein domains and mapping of 25 different mutations reported in our GLUT1DS patients (adapted from Galochkina *et al*., 2019) [23]. Novel variants identified in our cohort are reported in bold.

Among the GLUT1DS pediatric patients, 21 mutations occurred *de novo*, resulting in sporadic cases, while 6 were familial cases with clinically and genetically affected relatives, bringing the total number of individuals with *SLC2A1* variants to 39. The pedigrees of the 6 families with autosomal dominant transmission are reported in Fig. 2.

**Figure 2:**
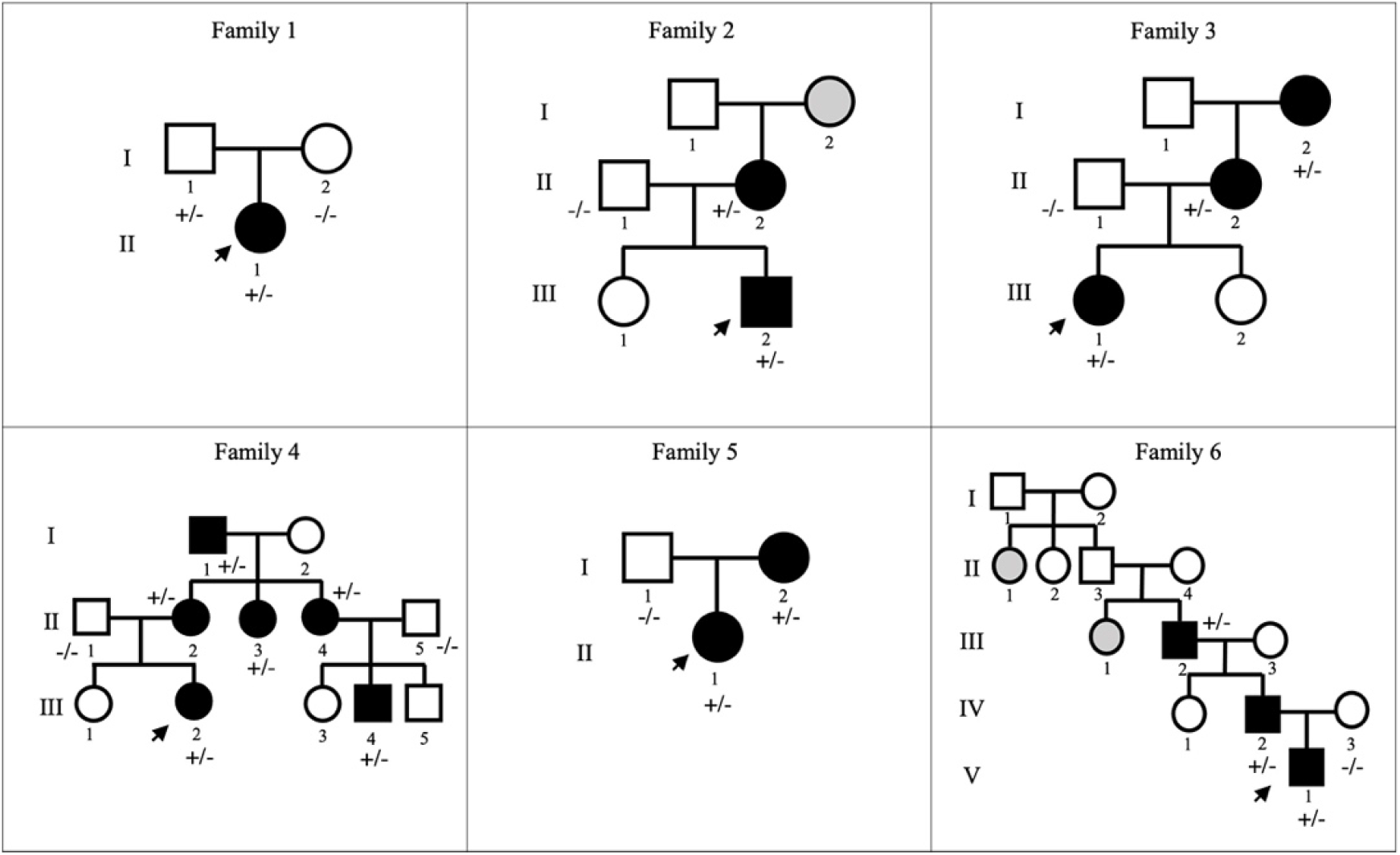
Pedigree of the familial cases with GLUT1DS. In family 2 and family 6, grey fills denote patients with clinical phenotype but unavailable for the study.

According to the ACMG criteria, 16 variants were classified as pathogenic, 5 variants reported as likely pathogenic and 4 variants of uncertain significance (VUS). Although these 4 variants (c.26C>T, c. 200T>C, c.275+3A>T and c.1363A>G) are classified as VUS, they were considered possibly causative based on the clinical phenotype of the patients, *in silico* predictions and/or family segregation.

### Genetic variations in Familial cases

We identified six families with autosomal dominant transmission of GLUT1DS.

#### Family 1

The proband (II-1; patient 1) was a 16-year-old girl presenting recurrent absence seizures since the age of 6 without movement disorder episodes. She was treated with different antiseizure medications, with no improvement in her condition. Her parents (I-1, I-2) are reported to be in good health.

Genetic analyses performed in the family trio revealed the presence of the heterozygous missense variant c.26C>T (p.Thr9Met) in exon 2 of the *SLC2A1* gene in the proband (II-1), inherited from the unaffected father (I-1). To determine the impact of this amino acid substitution in position 9, the pathogenic score was evaluated by nineteen different *in silico* predictors, and it was predicted to be damaging by seventeen of them (Table S1). This score is consistent with the amino acid change from a polar threonine to a non-polar methionine, in a highly conserved residue. The identified variant (rs1570601100) is classified as VUS in ClinVar (SCV000953099), reported as likely pathogenic in LOVD database and as a likely disease-causing mutation for GLUT1DS by HGMD Professional 2021.3. This same variant (c.26C>T) has been reported, for the first time, by Castellotti *et al*. in an individual affected with GLUT1DS and, as in our family 1, also in the asymptomatic father [29].

#### Family 2

The proband (III-2, patient 4) of this family was a 16-year-old boy suffering recurrent uncontrolled lower limb movement since the age of 7; free periods of six months were intercalated by weeks in which PED occurred every day (after intense exercise, or at the end of a stressful day). At the age of 2, the patient had two absence-like-episodes, not related to fever. The mother (II-2), a 49-year woman, suffered from recurrent episodes of paroxysmal exercise induced dyskinesia during adolescence (since 13 years old), then spontaneous resolution occurred in her early twenties. Moreover, the family reported seizures in maternal grandmother (I-2, not available for sequencing). Trio sequencing analyses identified the novel heterozygous missense variant c.200T>C (p.Leu67Pro) in exon 3 of the *SLC2A1* gene in the proband (III-2), inherited from the affected mother (II-2). This amino acid change from leucine to proline (p.Leu67Pro) affects a highly conserved on transmembrane helix 2 (TMH2), likely disrupting the domain’s α-helical architecture. The pathogenic score, indeed, was evaluated by nineteen different *in silico* predictors and it was predicted to be damaging by eighteen of them (Table S1). The *SLC2A1* variant identified in this family is not present neither in ClinVar nor in HGMD Professional 2021.3.

#### Family 3

The proband (III-1, patient 9) was a 9-year-old girl with recurrent generalized tonic-clonic seizures and absence seizure since the age of 3. With growth, during childhood, she suffered from dyskinesia triggered by exercise or fatigue to the lower limbs. EEG showed slow background activity generalized and focal discharges. The mother (II-2), a 41-year-old woman, affected by generalized tonic-clonic seizures, since the age of 2. She suffered from paroxysmal exercise induced dyskinesia, worsened with growth, mainly in stressful periods. Those movements affected, with the same severity and same frequency, both arms and legs. The maternal grandmother (I-2) was a 73-year woman with several generalized tonic-clonic seizures since the age of 17 maintained under control with a combination of different antiepileptic drugs. Since the age of 8, she also developed PED, which has worsened with growth. The lower limbs, especially her feet, were the most affected. In the last years, these movement disorders became rare, occurring only at the end of intense exercise-fatigue. Sequencing analysis allowed the identification of the heterozygous missense variant c.388G>A (p.Gly130Ser) in exon 4 of the *SLC2A1* gene in the proband (III-1), in the mother (II-2) and in the grandmother (I-2). To determine the impact of this amino acid change, the pathogenic score was evaluated by nineteen different *in silico* predictors, and it was predicted to be damaging by all of them (Table S1). The Gly130Ser mutation occurs at the narrow interface between TMH4 and TMH2, replacing a glycine, which lacks a side chain, with a cysteine whose side chain could sterically clash with TMH2, thereby destabilizing the helical packing of the transporter (Fig. S1b).

The identified variant (rs80359819) is classified as pathogenic in ClinVar (RCV000770978), as likely pathogenic in LOVD database, and as a disease-causing mutation for GLUT1DS by HGMD Professional 2021.3. It has been previously identified in GLUT1DS patients [24].

#### Family 4

The proband (III-2, patient 11) of this family was a 17-year-old girl with no seizure history, but recurrent paroxysmal exercise induced dyskinesia since the age of 6. This movement disorder was characterized by worsening due to fatigue and fasting. The mother (II-2), a 38-year woman, affected by generalized tonic-clonic seizure and absence seizure from the age one, with seizures every two-three days. She was on therapy from the age of one, with a combination of different antiepileptic drugs, with poor outcome. She was also affected by paroxysmal exercise induced dyskinesia since the age 25, with legs more affected than arms. Also a stroke-like episode occurred, with paresis of the upper limb and loss of language functions for 12-24 hours. The aunt (II-3) was affected by absence seizures since the age of 6. She has not developed PED episodes during her life. The aunt (II-4) of the proband (III-2) was a 34-year-old woman who suffered of absence seizures since the age of 6, once every two-three weeks. Since the age of 20, she has experienced paroxysmal exercise induced dyskinesia episodes. When she was 31 years old, she had two stroke-like episodes with left hemiparesis. She has 3 sons, one (III-4, a 7-year-old boy) affected by generalized epileptic seizure, mild intellectual disability and attention-deficit/hyperactivity disorder (ADHD); the other one by ADHD, and the last one, apparently, in healthy conditions. The maternal grandfather (I-1) was a 71-year-old man; he had recurrent migraines during adolescence, with dizziness just at the end of a physical effort, and PED. No other pathology was referred.

The heterozygous missense variant c.493G>A (p.Val165Ile) in exon 4 of the *SLC2A1* gene was identified in all family members showing epilepsy and/or PED episodes. This amino acid change (p.Val165Ile) affects a highly conserved residue whose position in TMH5 is crucial for glucose transport. The pathogenic score, indeed, was evaluated by nineteen different *in silico* predictors, and it was predicted to be damaging by fifteen of them (Table S1). The identified variant (rs1057520545) is classified as pathogenic in ClinVar (SCV002247281) and LOVD database, and as a disease-causing mutation for GLUT1-DS by HGMD Professional 2021.3. This same variant (c.493G>A) has been previously observed in individuals with GLUT1DS [9, 30].

#### Family 5

The proband (II-1, patient 16) was a 7-year-old girl presenting recurrent absence seizures since the age of 6 months. Moreover, since the age of 2, she is affected by exercise-induced dyskinesia. The mother (I-2), a 40-year woman, affected by seizure since childhood. She suffered from paroxysmal exercise induced dyskinesia episodes, mainly in stressful periods. She had a stroke episode. NGS analysis performed in the family trio showed the heterozygous missense variant c.884C>T (p.Thr295Met) in exon 8 of the *SLC2A1* gene in the proband (II-1), inherited from the affected mother (I-2). To determine the impact of this amino acid substitution in position 295, the pathogenic score was evaluated by nineteen different *in silico* predictors, and it was predicted to be damaging by eighteen of them (Table S1). This score is consistent with the amino acid substitution from a polar threonine to a non-polar methionine, in a highly conserved domain (TMH7) comprising the GLUT1 cavity. This is also supported by functional studies on this variant [31]; in particular, the authors showed that this amino acid change (p.Thr295Met) specifically alters the GLUT1 conformation and affects the glucose uptake. The rs80359823 variant is classified as pathogenic in ClinVar (SCV001577959) and LOVD database, and as a disease-causing mutation for GLUT1DS by HGMD Professional 2021.3, as it has been previously identified in GLUT1DS patients [24]. This same *SLC2A1* variant (c.884C>T) was also showed in patient 17, where both parents had been tested, as a *de novo* mutation, resulting in a sporadic case.

#### Family 6

The proband (V-1, patient 25) was a 3-years-old boy. Since the age of 4 months, he suffered from several partial seizures, characterized impaired awareness, cyanosis, apnea and tonic spasms. Frequent events in the same day (9-10 times in a day) were interrupted by seizure-free periods. The duration of each episode ranged from 10 seconds up to 30 - 40 seconds. The father (IV-1) suffered of paroxysmal exercise induced dyskinesia since the age of 10. The most affected limbs were the legs, and the episode duration was related to the exercise intensity. During adolescence those episodes worsened in terms of frequency and intensity, occurring daily and in his twenties, these episodes became less frequent. The paternal grandfather (III-2) showed recurrent PED, right lower limb was mainly affected (rarely arms were involved). It started at the age of 12, with increased intensity and frequency up to 20 years old, mainly 1-2 times in a month. These abnormal movements were mainly associated with stressful periods or right after physical exercise. Nowadays, the only remaining symptom are the migraine episodes that has been occurring throughout his life, starting since adolescence. Symptoms also occurred in other family members: sister of patient III-2 (III-1) was affected by recurrent and severe seizures, and anxiety disorder; the aunt of patient III-1 and III-2 (II-1) was affected by seizure and movement disorder (II-1, III-1 not available for sequencing).

The novel heterozygous missense variant c.1363A>G (p.Thr455Ala) in exon 10 of the *SLC2A1* gene was identified in the proband (IIIII-1), in the father (IIII-2) and in the grandfather (III-2). The *SLC2A1* variant identified in this family is neither classified in ClinVar nor in HGMD Professional 2021.3. To determine the impact of this amino acid substitution (p.Thr455Ala), the pathogenic score was evaluated by nineteen different *in silico* predictors, and it was predicted to be damaging by eighteen of them (Table S1). This score is consistent with the amino acid change in position 455 from a polar threonine to a non-polar alanine, in a highly conserved residue of IC5 involved in the GLUT1 cavity.

### Genetic variations in Sporadic cases

Whereas in all familial cases (100%) we detected missense mutations, only 52,4% (11 out of 21) of sporadic cases had missense variants; in the remaining 10 patients with *de novo* mutations, we identified frameshift (28,6%) and nonsense, splice site, in-frame and noncoding variants (19%). We found 14 pathogenic variants, 6 likely pathogenic and one VUS, according to ACMG criteria.

To note, the variant classified as uncertain significance was a heterozygous *de novo* noncoding mutation c.275+3A>T detected in patient 6. It has been previously identified in GLUT1DS patient by Graziola *et al*. [32], and considered causative, as in our case, based on clinical presentation of the proband. Further *in silico* analysis on Genomnis HSF suggested that this variant (c.275+3A>T) most probably affected the splicing process, leading to alteration of the 5’ splice site in intron 3.

### Novel SLC2A1 variants and their crystallographic structure

Out of a total of 25 identified variants, 40% have never been reported before (8 *de novo* and 2 inherited), including one splice site (10%), 4 frame shift (40%), 5 missense (50%).

The novel heterozygous *de novo* splice site variant c.114+1G>A identified in patient 3 affected the donor splice site of intron 2, possibly leading to exon 2 skipping. This hypothesis was confirmed by the *in silico* analyses of the splicing process of the alternative transcript. Splicing prediction tools (e.g. Fruit Fly Splice Predictor, ESEfinder 3.0, Genomnis HSF), indeed, suggested that the novel variant c.114+1G>A causes the loss of the 5’ splice site in intron 2 and the consequent skipping of exon 2, thus leading to exon 1-3 junction (Fig. 3). In particular, the in-frame loss of exon 2 causes the completely loss of the first transmembrane helix (TMH1) of the GLUT1 (predicted by protein topology THMM *in silico* tool), which is highly involved in glucose uptake, probably due to its important role for the protein state transition [23].

**Figure 3:**
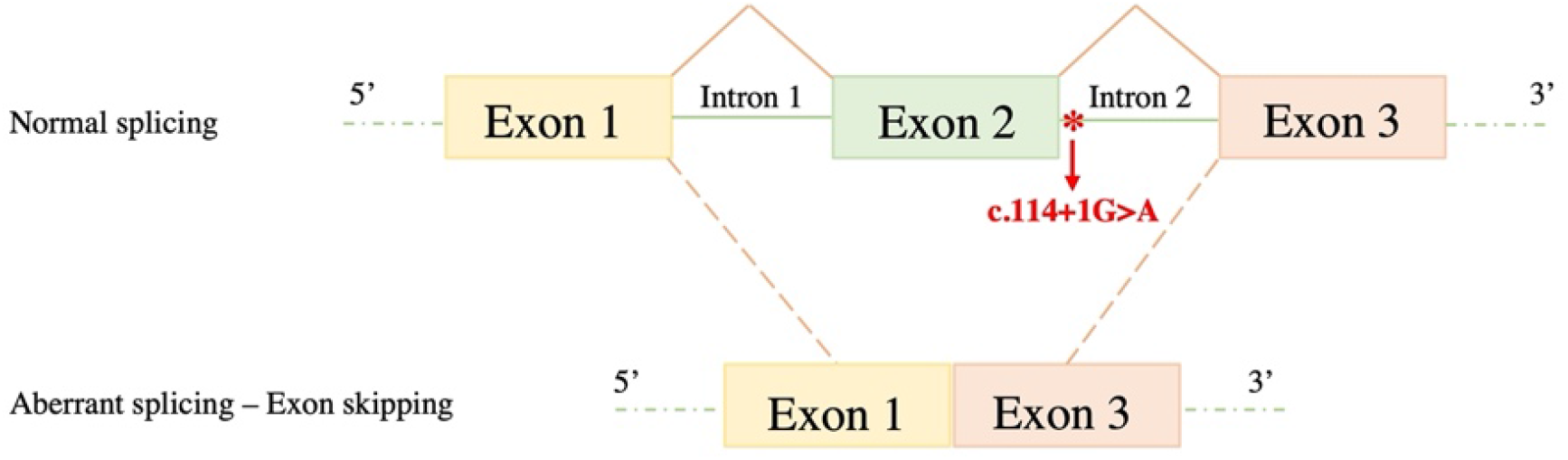
Schematic illustration of normal splicing (up) and aberrant splice event (above), which results in exon 2 skipping. The red asterisk indicates the novel heterozygous *de novo* c.114+1G>A variant in intron 2.

The novel frameshift mutations identified in our patients cause the introduction of a PTC, predicted to lead to the synthesis of non-functional truncated protein, or more likely to NMD. It is likely that these mutations result in 50% loss of GLUT1 protein and thus, lead to severe impairment of glucose transport into brain. These molecular findings were also supported by both biochemical and clinical data of patients 7,14, 18 and 27: the CSF/blood glucose ratios were 0,36, 0,37, 031 and 0,37 mg/dL, respectively; all patients had the early-onset severe phenotype in combination with intellectual disability and/or movement disorders.

The amino acid changes in the novel missense variants identified in this study were evaluated by nineteen different *in silico* predictors, and they were predicted to be damaging by most of them (Table S1–S2). Additionally, in order to rationalize how these novel disease-related missense mutations may impact the structure and function of human GLUT1 (hGLUT1), we mapped them on its X-ray crystallographic structure in complex with Nonyl-β-D-Glucoside [33, 34] (PDB ID: 6THA) (Fig. 4a). The latter is a detergent bearing a glucoside group which binds to the same hGLUT1 pocket that in other structures of hGLUT1 homologues is occupied by the native ligand glucose (for example in hGLUT3 structure, PDB ID: 4ZW9 [35] and the *E. coli* homolog of GLUT1-4, XylE, PDB ID: 4GBZ [36]). Similarly, in another equivalent reported structure of hGLUT1, the same pocket is occupied by the cytochalasin B inhibitor (PDB ID: 5EQI) [37]. As follows, we discuss the potential significance of each newly identified variant in the context of their chemical environment and position in key secondary structure elements of hGLUT1 (Fig. 4 b-e). The Leu67Pro mutation, located on TMH2, may cause a kink of the α-helix due to the rigidity of the cyclic pyrrolidine side chain of proline (Fig. 4b). Despite not being directly involved in ligand binding, this mutation could affect the overall protein helix bundle flexibility and therefore the glucose transport efficiency. The Gly75Glu variant, also found in TMH2, corresponds to the replacement of a small hydrophobic glycine residue with a negatively charged glutamic acid on the membrane-facing side of TMH2 (Fig. 4b). Here, the mutation could in principle affect not only the folding, but also the localization of hGLUT1 in the phospholipidic membrane. Gln283Lys is found right in the internal cavity of the hGLUT1 transporter where glucose transits (Fig. 4c). Here, the WT residue Gln283 forms a hydrogen bond network with Asn288, likely involved in stabilizing the glucose head group of the Nonyl-β-D-Glucoside, mimicking the natural ligand. The replacement with a longer and charged residue as lysine may likely disrupt this highly specific binding configuration, probably modifying the glucose passage through the channel. Moving to a different region of the transporter, the Ala377Pro mutation located on the transmembrane helix 10 (TMH10) could potentially have a similar destabilizing effect as the Leu67Pro mutation in TMH2 (Fig. 4d). The Thr455Ala mutation is located within the C-terminal portion of hGLUT1 in a flexible hinge after TMH12 (Fig. 4e). Interestingly, Thr455 establishes a network of interactions with Glu393 on TMH10 and with Arg333 on TMH9, holding the three C-terminal α-helices together. These may also explain why Thr455 is the last semi-rigid and visible residue in the crystallographic model. The pathogenicity of the Thr455 to alanine mutation could probably be related to the disruption of this C-terminal “lock” network with a detrimental increase on the C-terminal domain flexibility and possibly on the dynamics of ligand transport.LA 377

**Figure 4:**
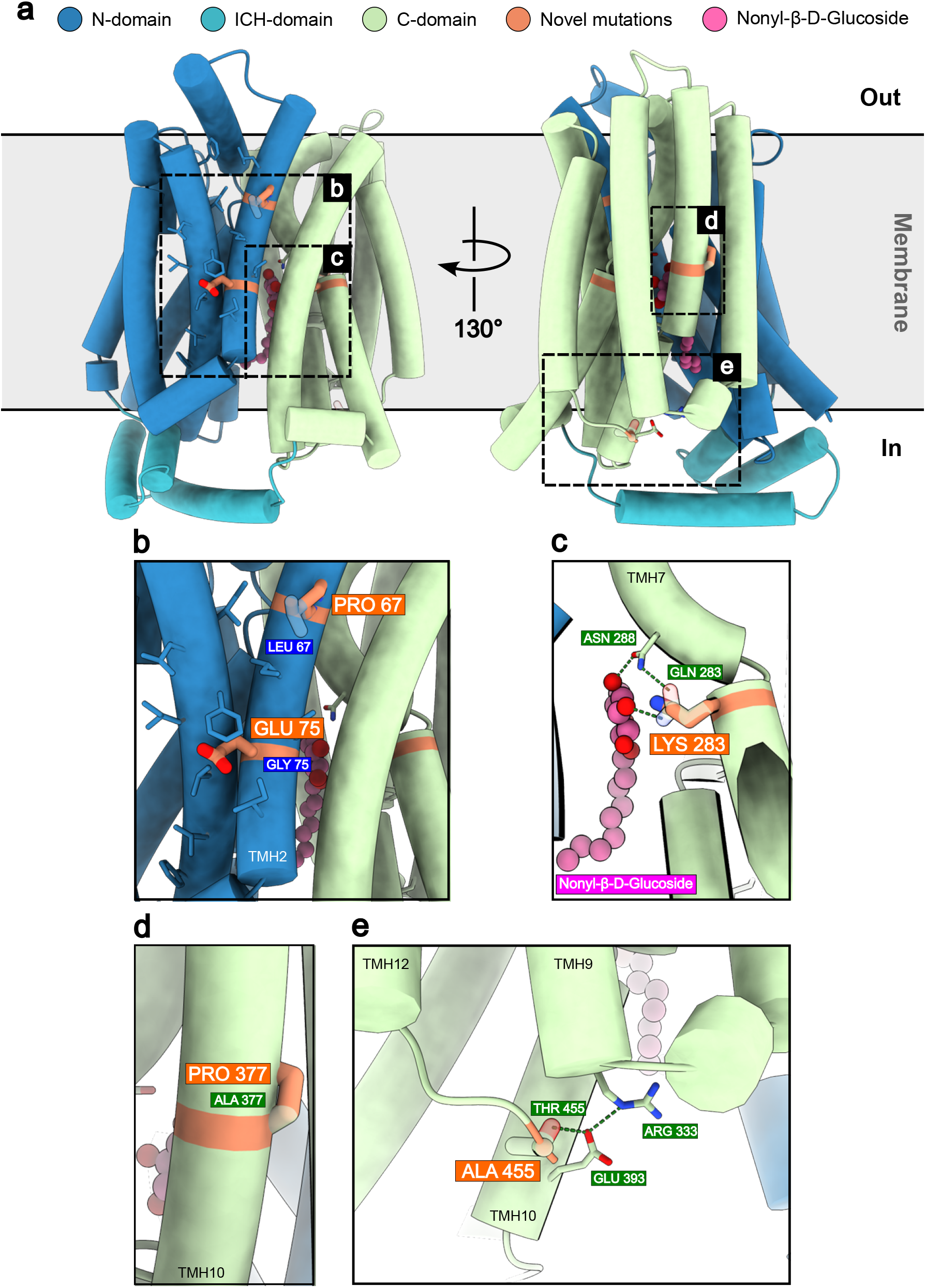
Novel disease-related mutations mapped on the structure of GLUT1. **a**, X-ray structure of GLUT1 (PDB ID: 6THA) bound to Nonyl-β-D-Glucoside via its glucose head group. Novel disease-related mutations identified in this study are depicted in orange and superimposed on their corresponding wild-type residues. **b**, Mutations Leu67Pro and Gly75Glu on transmembrane helix 2 (TMH2) could disrupt the α-helical architecture and the hydrophobicity of the region, respectively. **c**, Gln283Lys could hamper the hydrogen bond network that stabilizes glucose binding. **d**, Ala377Pro could destabilize the α-helical architecture of TMH10. **e**, Thr455Ala potentially breaks the hydrogen bond network between residues on TMH9, TMH10 and TMH12, possibly reducing the stability and rigidity of the C-terminal domain of GLUT1.

### Genotype, phenotype and biochemical correlations

Given the clinical variability and the wide spectrum of heterozygous mutations (including missense, frameshift, nonsense, and splice site variants) observed in our study cohort, we investigated possible associations between phenotype, genotype, and biochemical data.

The early-onset classical phenotype was seen in both patients with missense mutations (59%; n = 10) and patients with splice site and frameshift variants (41%; n = 7). All the patients with splice site and frameshift variants had the early-onset classical features, whereas 41% (n = 7) of the patients with missense mutations had the late-onset or non-classical phenotype (Fig. 5). Patients with identical missense mutations (16, 17 and 24, 25) were heterogeneous in clinical manifestations.

**Figure 5:**
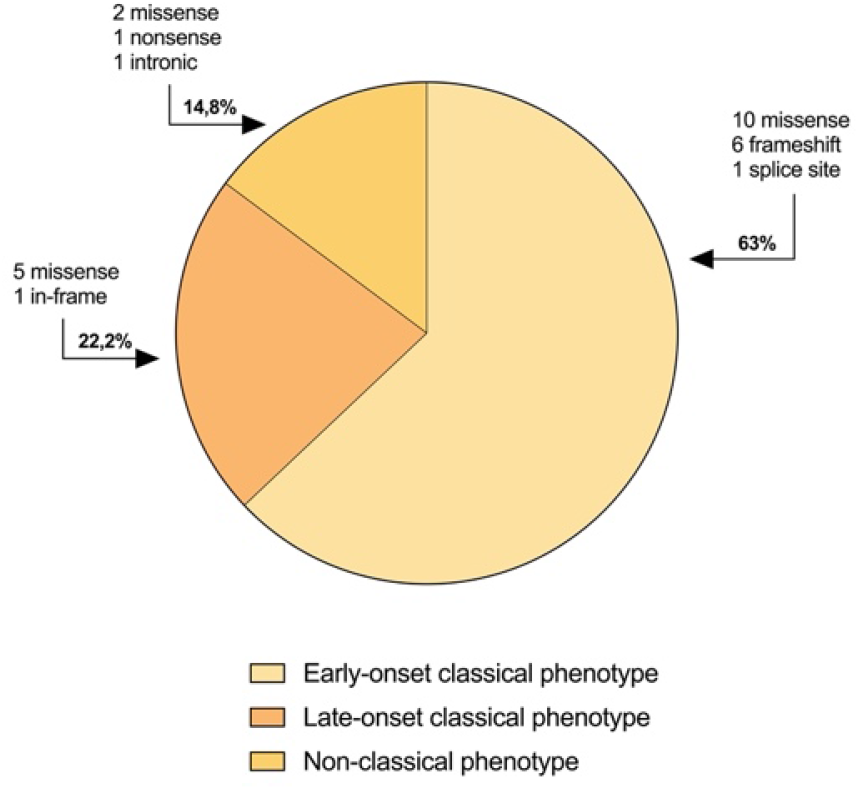
Distribution of the 27 *SLC2A1* variants observed in our cohort according to the different phenotype.

Patients with the late-onset classical phenotype had a higher CSF/blood glucose ratio (mean 0.44; SD 0.07) than patients with the early-onset classical phenotype (mean 0.37; SD 0.05) and patients with non-classical clinical features with movement disorders without epilepsy (mean 0.35; SD 0.05) (Fig. 6a).

**Figure 6:**
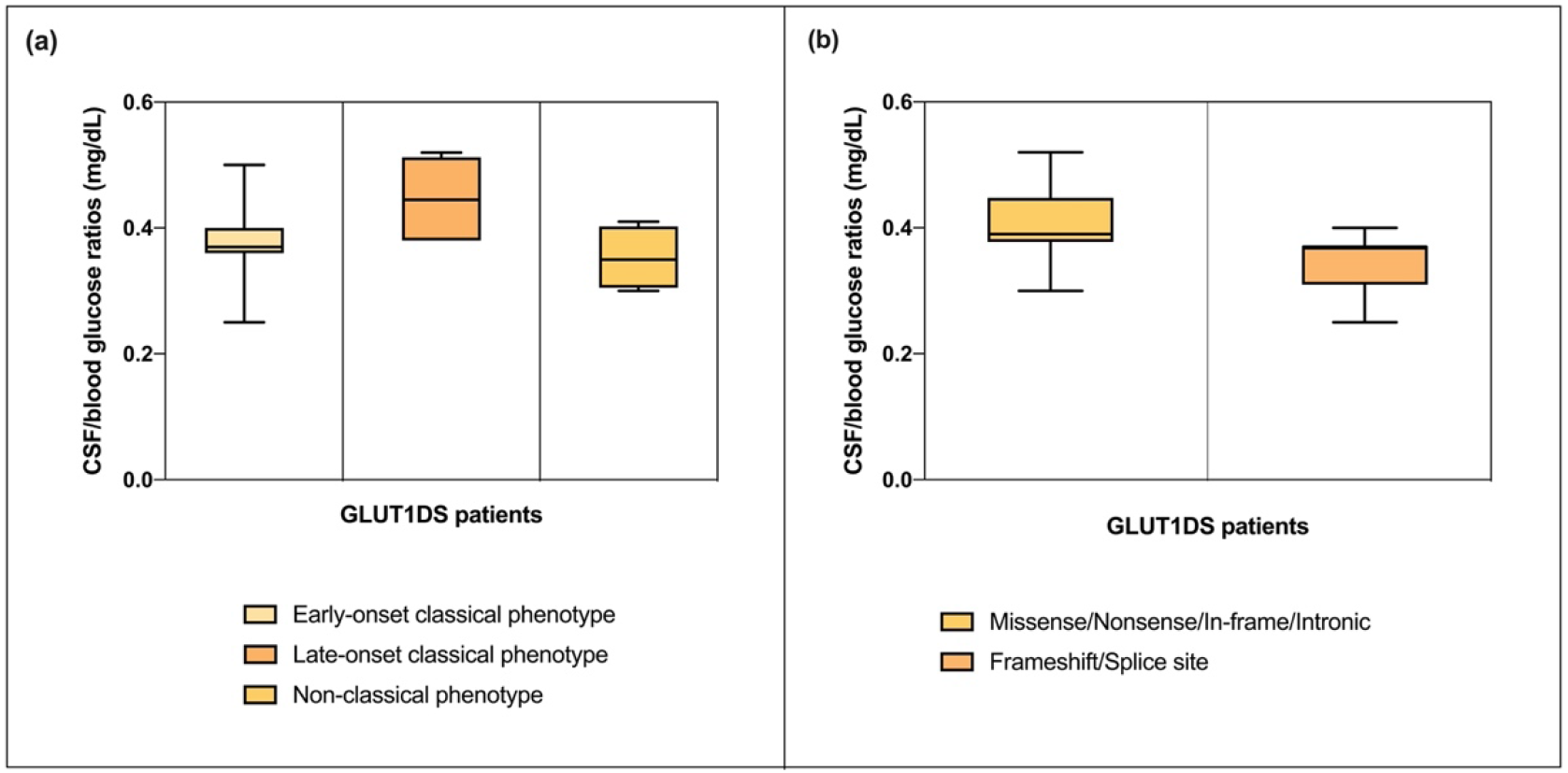
Genotype, phenotype and biochemical correlations. **a**, Boxplot shows CSF/blood glucose ratios with three different phenotypes. **b**, Boxplot displays CSF/blood glucose ratios with different type of variants.

The CSF/blood glucose values were lower in patients with splice site and frameshift mutations (mean 0.34; SD 0.05) than in individuals with missense mutations (mean 0.40; SD 0.05) (Fig. 6b).

## Discussion

For the first time, we reported the molecular data of 27 Italian children with a clinical suspect of GLUT1DS; specifically, we performed *SLC2A1* sequencing analysis, investigated possible correlations between phenotype, genotype and biochemical data and then highlighted novel disease-related mutations and a potential mutational hotspot in exon 4 of the *SLC2A1* gene. In our cohort we have identified 17 patients (62,9%) with the early-onset classical phenotype, 5 patients (18,5%) with the late-onset classical phenotype, while in the remaining 4 patients (14,8%) we observed a non-classical phenotype of movement disorders without epilepsy. This latter phenotype is considered uncommon and has only been described in a small number of patients [12,14]. We have, however, identified in patient 19 only minimal symptoms such as intellectual disability and microcephaly, suggesting a wide clinical spectrum of GLUT1 deficiency syndrome.

CSF/blood glucose ratios were below 0.52 mg/dL in all our patients, and below 0.37 mg/dL in patients with splice site and frameshift mutations (25,9%), as showed in Fig. 6b, and thus suggesting that genotype could be correlated with CSF parameters. All these mutations, resulting in 50% loss of GLUT1 activity, are identified in patients with the early-onset classical phenotype in combination with intellectual disability and/or movement disorders (patients 2, 3, 7, 14, 18, 21, 27). This highlights that low CSF/blood glucose ratios are most likely related to a more severe phenotype, as well as with splice site and frameshift variants.

Our molecular findings showed important differences between sporadic and familial cases in the type of mutations, and thus in the severity of the clinical manifestations. Heterozygous missense variants detected in all familial cases, as is now well established [38], were associated with mild to moderate forms of the disease with broad phenotypic variability: the early-onset classical phenotype was observed in patients 9 (family 3), 16 (family 5) and 25 (family 6), the late-onset was noted only in patient 1 (family 1), whereas patients 4 (family 2) and 11 (family 4) presented non-classical features with movement disorders without epilepsy. Furthermore, we observed wide clinical heterogeneity even within the same family [39]. Patients with the same mutation (c.493G>A) in family 4 displayed a heterogenous range of type and severity of GLUT1DS phenotype. In family 1 the identified c.26C>T variant is present in the affected proband (patient 1) and in the unaffected father. This phenotypic diversity within families with autosomal dominant transmission of GLUT1DS suggests a modulating effect of DNA regulatory elements of the *SLC2A1* gene that may modulate the expression level of the GLUT1 protein. This assumption emphasizes the importance of further clinical and genetic/epigenetics investigation of GLUT1DS families.

Overall, 40% of the identified variants in our *SLC2A1* cohort (10 out of 25) had never been reported before, including missense, frameshift and splice site variants, broadening the genotypic spectrum heterogeneity found in the *SLC2A1* gene. Although two of these (c.200T>C; c.1363A>G) were classified as VUS according to the ACMG criteria, X-ray structure analyses of GLUT1 strongly suggested the potential pathogenic nature of these novel variants (Fig. 4), providing additional evidence to overcome the challenging VUS classification. Instead, the pathogenicity of the two VUS identified in patient 1 and patient 6 was supported by literature data [29,32].

A significant fraction (6 out of 25; 24%) of the identified variants is located in a vulnerable region of the GLUT1 protein that involves TMH4 and TMH5 domains encoded by exon 4, as showed in Fig. 1b and represented on the same reference hGLUT1 structure in Fig. S1. TMH4 and TMH5 include mutations at amino acid residues Leu124, Arg126, Gly130, Arg153, Val165 and Val166, which are located around GLUT1 central channel and involved in the interaction with the glucose. During the protein state transition for the glucose uptake, many residues, including Arg126 on TMH4, form strong hydrogen bonds with the glucose molecules. Of note, the amino acid position 126 is exposed at the extracellular part of the GLUT1 protein and a positively charged arginine facilitates glucose transport, resulting in a glucose binding site residue [23]. Thus, the amino acid change from a positively charged arginine to a shorter polar side chain of cysteine (p.Arg126Cys) observed in patient 8, could affect the binding affinity of GLUT1. At the same time, the disruption of the salt bridge at the intracellular part of the transporter between TMH5, TMH10 and IC3 (Arg153, Glu397 and Glu247) due to the Arg153Cys mutation on TMH5 (detected in patient 10) could lead to the dramatical decrease of the glucose uptake [23]. These molecular findings are also supported by both biochemical and clinical data of patients 8, 9 and 10: the CSF/blood glucose ratios are 0.34, 0.39 and 0.39 mg/dL, respectively. Also stroke episodes occurred in patients 8 and 10, suggesting a critical functional disturbance associated with structural alterations in TMH4 and TMH5 domains of the GLUT1 transporter.

In summary, these molecular data confirmed also in our cohort the mutational hotspot in exon 4 of the *SLC2A1* gene observed by Pascual *et al*. [28], suggesting that structural alterations in TMH4 and TMH5 domains of the GLUT1 transporter could affect the glucose uptake into the brain. Additionally, our discovery of the ten novel disease-related variants and their X-ray structure analyses broadens the genotypic spectrum heterogeneity found in the *SLC2A1* gene, suggesting the pathogenic effects of these identified mutations. Lastly, the wide clinical and genetic heterogeneity observed in our GLUT1DS cohort allowed possible correlations between mutation type and clinical and biochemical data. This analysis enabled to delineate that splice site and frameshift variants are related to a more severe phenotype characterized by early-onset classical phenotype and low CSF parameters.

## Materials and methods

### Study population

We enrolled 27 Italian pediatric patients who were referred to the *SLC2A1* gene analysis by the physicians of Buzzi Children’s Hospital in Milan for a clinically suspect of GLUT1DS. For all patients we sequenced trios (proband and parents), and when available other family members (i.e. family 3, family 4, and family 6). The probands include 17 females and 10 males with ages ranging from 5 months to 19 years (average 6 years) at the time of recruitment.

This study was approved by the ethics committee of Area 1 of Milan (2021/ST/004) and written informed consent was obtained from all pediatric patients and their family.

### CSF biochemical analysis

In all patients, cerebrospinal fluid (CSF) was collected through the spinal tap procedure. Then, the quantification of glucose in CSF was performed using the Alinity c Glucose enzymatic assay (Abbott Laboratories).

### Mutation analysis of the SLC2A1 gene

Genomic DNA from probands and relatives were extracted from peripheral blood leukocytes (PBLs) using a semi-automated method Maxwell System DNA Purification (Promega, Madison, WI, USA), according to the manufacturer’s instructions. The genetic analyses have been performed Next Generation Sequencing (NGS) panels.

NGS analyses were performed using an Agilent custom SureSelect panel (Agilent, Santa Clara, CA) and libraries were sequenced on the Illumina NextSeq 550 platform (Illumina, San Diego, CA). Sequences were aligned to the reference genome (GRCh38), and variants were called through the Sentieon (Sentieon Inc, San José, CA, USA) and annotated using VarSeq (Golden Helix, Bozeman, MT, USA). Reads with low quality calling score and coverage <10, reads falling in segmental duplicated regions (SuperDups>0.9), and common SNPs (Minor allele Frequency (MAF) >1%) were filtered and removed.

Sanger sequencing was used to confirm the NGS variations found in patients or to define variations in parents or other family members. We used a standard protocol to amplify all 10 exons of the *SLC2A1* gene using primers located in adjacent intronic regions on gDNA by polymerase chain reaction (PCR). All amplicons were screened for sequence variations by direct sequencing using Big-Dye Terminator v3.1 sequencing kit (Applied Biosystems, Foster City, CA, USA) and ABI 3130 Genetic Analyzer (Applied Biosystems, Foster City, CA, USA). Each fragment was sequenced on both strands. The alignment to the reference sequence (NM_006516) was performed using Sequencher 4.8 software (Gene Codes Corporation, Ann Arbor, MI, USA).

The identified variants in the *SLC2A1* gene were interpreted according to the American College of Medical Genetics and Genomics (ACMG) guidelines [40] with the support of several tools such as Varsome, Franklin by Genoox (https://franklin.genoox.com) and locus specific-variant databases (including HGMD and ClinVar). Several *in silico* prediction tools (e.g. SIFT, PolyPhen2, MutationTaster) were used to assess pathogenicity scores; in particular, we used the MutationTaster tool (https://www.mutationtaster.org) to define the position of premature termination codons (PTCs) compared to the canonical ones and to predict the effect of nonsense-mediated decay (NMD) according to the rules governing this surveillance process [41].

GLUT1 structural analysis related to patient variations in Fig. 4 and Fig. S1 were performed with UCSF ChimeraX [42].

## Acknowledgments

The authors acknowledge the probands and their families for participating in this study and giving consent to publish their data.

---

**Table S1:**
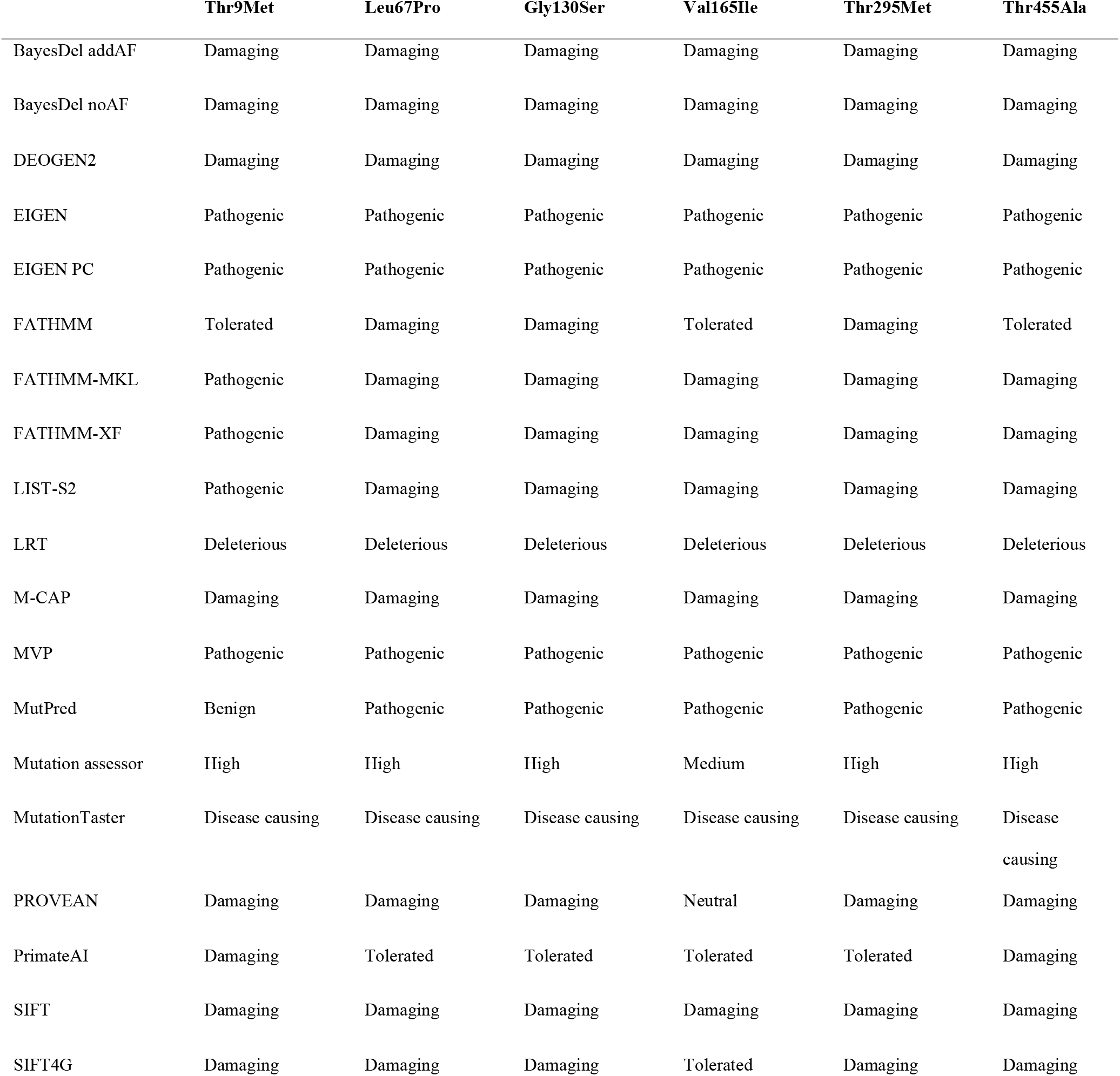
Prediction of pathogenicity of the identified inherited missense GLUT1 variant by *in silico* tools.

**Table S2:** Prediction of pathogenicity of the identified novel *de novo* missense GLUT1 variant by *in silico*

|  | Gly75Glu | Gln283Lys | Ala377Pro |
| --- | --- | --- | --- |
| BayesDel addAF | Damaging | Damaging | Damaging |
| BayesDel noAF | Damaging | Damaging | Damaging |
| DEOGEN2 | Damaging | Damaging | Damaging |
| EIGEN | Pathogenic | Pathogenic | Pathogenic |
| EIGEN PC | Pathogenic | Pathogenic | Pathogenic |
| FATHMM | Damaging | Tolerated | Damaging |
| FATHMM-MKL | Damaging | Damaging | Damaging |
| FATHMM-XF | Damaging | Damaging | Damaging |
| LIST-S2 | Damaging | Damaging | Damaging |
| LRT | Deleterious | Deleterious | Deleterious |
| M-CAP | Damaging | Damaging | Damaging |
| MVP | Pathogenic | Pathogenic | Pathogenic |
| MutPred | Pathogenic | Pathogenic | Pathogenic |
| Mutation assessor | High | High | High |
| MutationTaster | Disease causing | Disease causing | Disease causing |
| PROVEAN | Damaging | Damaging | Damaging |
| PrimateAI | Damaging | Damaging | Damaging |
| SIFT | Damaging | Damaging | Damaging |
| SIFT4G | Damaging | Damaging | Tolerated |

**Figure S1:**
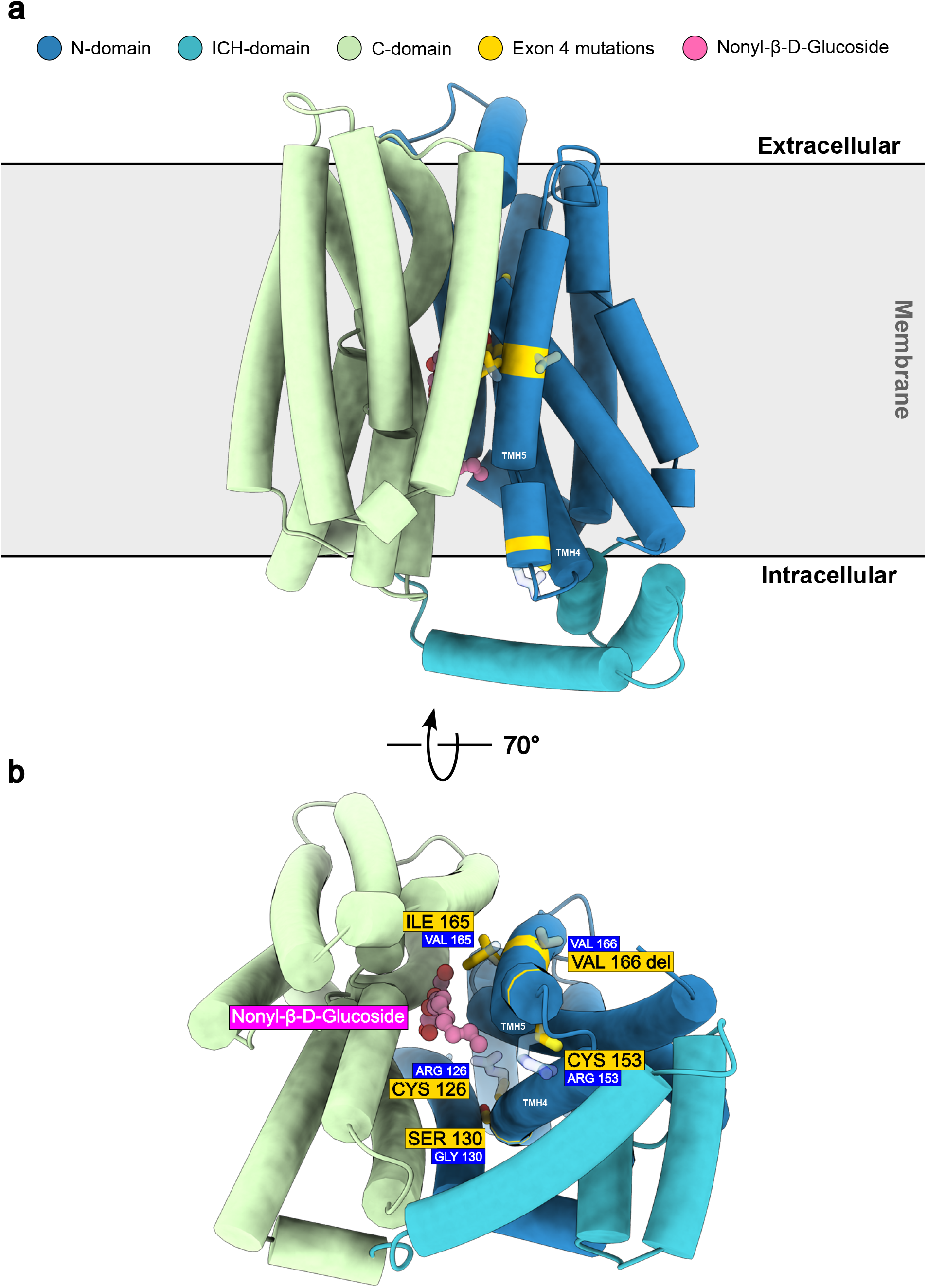
Exon 4 of the human *SLC2A1* gene is a hotspot for disease-related mutations. **a**, X-ray structure of GLUT1 (PDB ID: 6THA) bound to Nonyl-β-D-Glucoside via its glucose head group. Disease-related mutations on Exon 4 targeting transmembrane helix 4 (TMH4) and 5 (TMH5) are depicted in yellow and superimposed on their corresponding wild-type residues. **b**, Intracellular view of hGLUT1. The disease-related mutations on Exon 4 target residues of transmembrane helix 4 (TMH4) and 5 (TMH5) of hGLUT1. The Arg126Cys and Arg153Cys mutations replace the large positively charged side chain of arginine with the shorter polar side chain of cysteine, with several potential detrimental consequences on hGLUT1 structure stability. The Gly130Cys mutation occurs at the narrow interface between TMH4 and TMH2, replacing a glycine, which lacks a side chain, with a cysteine whose side chain could sterically clash with TMH2, thereby destabilizing the helical packing of the transporter. The Val165Ile mutation changes a smaller valine to a larger isoleucine at the interface between two helices and relatively close to the substrate binding cavity. This mutation could potentially affect the helical packing and maybe to a lesser extent the interaction with glucose. Although the mutation Val166del is a single aminoacidic deletion (in frame), it shifts the register of TMH5 residues, misplacing the outward-facing hydrophobic residues and the inward-facing polar residues with potentially devastating consequences on protein folding and function.**Table S1:** Prediction of pathogenicity of the identified inherited missense GLUT1 variant by *in silico* tools.

